# Many circulating indole and phenol metabolites are host derived

**DOI:** 10.1101/2025.10.05.680522

**Authors:** Jenna E. AbuSalim, Kellen Olszewski, Salma Youssef, Craig J. Hunter, Michael R. MacArthur, Alessa Henneberg, Rolf-Peter Ryseck, Christiane A. Opitz, Mohamed S. Donia, Joshua D. Rabinowitz

## Abstract

Indole and phenol metabolites are typically thought to be products of bacterial digestion of tryptophan (indoles) and phenylalanine or tyrosine (phenols). Interest in controlling gut microbial production of these metabolites has continually grown as they have important physiological impacts, with indoles agonizing AhR signaling, and phenols being associated with healthy body weight. While there is a growing wealth of research into which bacteria produce these metabolites, host contribution to their circulating pools has not been adequately characterized. Here, through stable isotope tracing in cell culture and mice, we show that mammalian cells can make aryl-pyruvates, -lactates, -acetates, and -carboxylic acids. Levels of these metabolites in mice and human patients are insensitive to perturbations of the microbiome. In contrast, bacterial metabolism is required to synthesize aryl-propionates and free indole, phenol, and cresol. Overall, we show that host metabolism is a primary contributor to circulating indole and phenol metabolite pools.

## Main

Indole and phenol metabolites have major impacts on host physiology (1–3). Indole metabolites are ligands for the aryl hydrocarbon receptor, which is a key mediator of gut immune tolerance and epithelial homeostasis (2–4). Phenylalanine-derived phenol metabolites such as cinnamic acid correlate with healthy body weight and can affect glucose and lipid handling (1), while tyrosine-derived metabolites such as phenol sulfate or cresol sulfate are uremic toxins (5,6). Supplementation with phenylalanine or tryptophan metabolites shows promise for the treatment of various diseases. In rodent models, providing indole metabolites alleviates disease severity of inflammatory bowel disease (7–9) while supplementing with cinnamic acid has protective effects against obesity and non-alcoholic fatty liver disease (1,10).

As indole and phenol metabolites have important physiologic impacts, there is growing interest into identifying which bacteria are responsible for their production (11). A recent study showed that Clostridium species have a metabolic pathway that can reduce all aryl-amino acids down to their respective propionates (12). Other species such as *Lactobacillus johnsonii* have also been shown to produce indole-3-pyruvate and phenylpyruvate as well as downstream metabolites (3,13,14). Additionally, changes in indole and phenol metabolites concentrations are often correlated with changes in microbiome composition (2,15).

While this work highlights a potential role of gut microbial health in controlling indole and phenol metabolite concentrations, microbiome compositional changes are also known to affect host metabolism (16). Recently, it was shown that kynurenine, a tryptophan metabolite made in macrophages, can be induced when mice are colonized with *Achromobacter pulmonis* (17). Additionally, the gut microbiome can even affect glucose homeostasis through various mechanisms including by producing short chain fatty acids that can fuel hepatic gluconeogenesis (16,18). IL4i1, a monocyte specific enzyme, has the capacity to produce indole-3-pyruvate and phenylpyruvate, the first metabolites in the indole and phenol reductive synthesis pathways respectively (19–21). This suggests that there may be some contribution of host metabolism to circulating indole and potentially phenol pools. Additionally, recent work showed comparable levels of some indole metabolites between germ free and conventional mice (22). Whether the presence of these metabolites in germ-free mice reflects uptake from diet or host metabolism remains to be determined (7,10). Isotope tracing, such as infusing ^13^C-labeled substrates, provides a gold-standard method for determining metabolite sources (23).

Here, we systematically evaluate the sources of indole and phenol metabolites based on (i) concentrations in germ-free mice, (ii) production over 2 h from infused ^13^C-labeled amino acids, which feed host cells but only minimally feed the gut microbiome, (iii) production over 36 h from infused ^13^C-labeled amino acids, which feeds both the host and, via secreted host protein, also the microbiome, (iv) modulation of labeling from the long duration infusions by antibiotics, and (v) direct production from ^13^C-labeled amino acids by mammalian cells in culture. Through these studies, we trace mammalian cell metabolism of aryl amino acids to their respective aryl-pyruvates, -lactates, -acetates, and -carboxylic acids (but not aryl-propionates and free indole, phenol, and cresol). We find that the microbiome-derived, but not host-derived, indole and phenol metabolites decline in human patients treated with antibiotics. Thus, many important indole and phenol metabolites that were thought to be microbiome derived are primarily synthesized by the host.

## Results

### Germ free mice produce indole metabolites

We began by measuring indole metabolites in serum from germ-free (GF) or control (Ctrl, or specific pathogen free) mice (Fig. 1A). Standard chow (PicoLab Rodent Diet 20, 5053) has a high abundance of microbiome metabolites including many of the indole species of interest. Accordingly, to identify endogenous sources of these metabolites, all mice were fed a doubly irradiated purified diet (D11112201-1.5Vii, Research Diets). Compared to chow, this diet had significantly lower levels of the indole metabolites (Fig. S1A).

**Figure 1:**
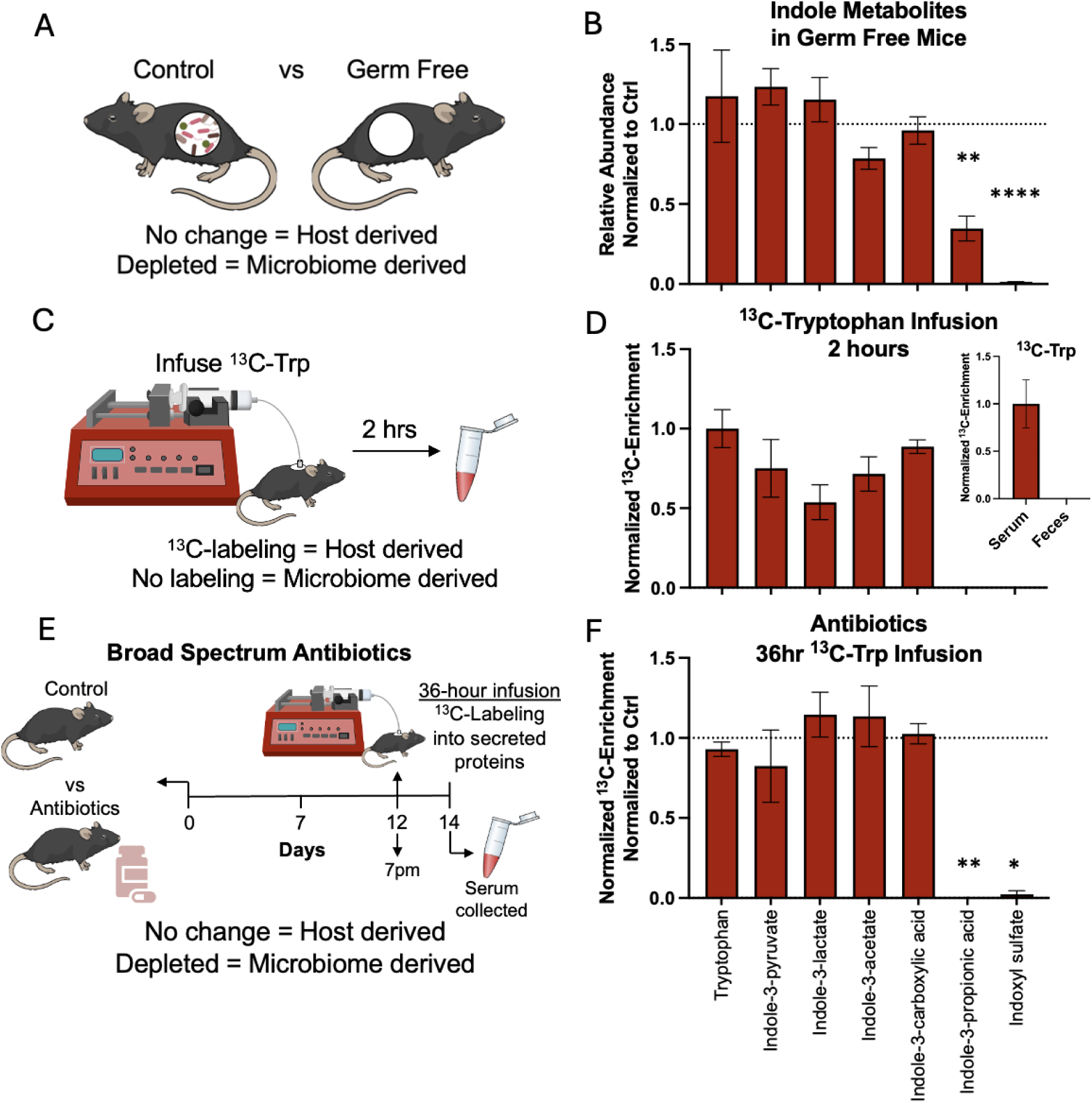
Many circulating indole metabolites are made by the host. (**A**) Scheme and (**B**) measurement of indole metabolites in serum of germ-free (GF) mice (n=5 female mice) compared to control (Ctrl) mice (n=4 female mice). While there is no microbial contribution in GF mice, circulating metabolite concentrations may be affected by direct dietary input of the metabolites. (**C**) Scheme describing the infusion of ^13^C-tryptophan for two hours and (**D**) ^13^C-enrichment of indole metabolites in serum normalized to serum ^13^C-tryptophan enrichment (n=3 male mice). Inset showing ^13^C-tyrptophan in serum and feces at 2 hours. (**E**) Scheme detailing a 14-day antibiotic treatment with ^13^C-tryptophan infused for the last 36 hours. (**F**) ^13^C-indole enrichment in antibiotic treated male mice (n=4) compared to enrichment of ^13^C-indole metabolites in control male mice (n=5). *p* values were calculated using a two-tailed *t*-test and notated as \**p*<0.05, \*\**p*<0.01, \*\*\**p*<0.001, and \*\*\*\**p*<0.0001.

Ten days after initiation of the purified diet, feces were collected to confirm sterility of the GF mice, and serum was collected and analyzed via liquid chromatography-mass spectrometry (LC-MS). Sterility was confirmed through anaerobic and aerobic cultures of dietary and fecal pellets. Some indole metabolites, such as indole-acrylic acid, were below the detection limit in serum from all mice. Others were readily detected. In serum from germ-free mice, indole-3-propionate was significantly depleted in comparison to control mice and indoxyl sulfate was completely absent. Other measured indole metabolites (indole-3-pyruvate, indole-3-acetate, indole-3-carboxylic acid, and indole-3-lactate), however, were not changed (Fig. 1B).

### Most circulating indole metabolites are host derived in vivo

To trace production of the indole metabolites, we infused minimally perturbative quantities of ^13^C-tryptophan into the systemic circulation of jugular-vein catheterized mice for 2 hours (Fig. 1C). As shown in prior work by our group (23), circulating amino acids do not readily cross into the gut lumen, making the infused ^13^C-tryptophan inaccessible to gut microbes. Consistent with this, there was no detectable ^13^C-tryptophan in the feces at 2 hours despite 20% circulating tryptophan labeling. There was also no labeling observed in circulating indole-3-propionate or indoxyl sulfate. The remaining indole metabolites, however, all were labeled to the same quantitative extent as tryptophan itself, supporting their being host derived and rapidly produced from circulating tryptophan (Fig. 1D). Based on the combination of these tracing data and the concentration data in germ-free mice, we assigned indole-3-pyruvate, indole-3-acetate, indole-3-carboxylic acid, and indole-3-lactate as host derived and indole-3-propionate or indoxyl sulfate as microbiome derived.

We next sought to observe ^13^C-labeing in the microbiome-derived indoles. For this purpose, we again infused ^13^C-tryptophan, but this time for a much longer duration (Fig. 1E). Previous work by our group showed bacteria in the gut microbiome utilized labeled amino acids that were infused for 36 h (23) as the infused amino acid can reach the gastrointestinal lumen through host secretion of proteins such as mucins. Fecal tryptophan labeling was indeed observed at 36 h (Fig. S1B). Serum measurements at 36 h revealed ^13^C-labeling into all indole metabolites, including indole-3-propionate and indoxyl sulfate, supporting a source of the microbiome-derived indoles being bacterial processing of tryptophan derived from secreted host protein (Fig. S1C). Side-by-side experiments in mice treated for 14 days with broad-spectrum antibiotics (ampicillin, neomycin, vancomycin, and metronidazole, ANVM) showed no difference in luminal tryptophan labeling as a function of antibiotics (Fig. S1B), but labeling of indole-3-propionate and indoxyl sulfate was largely ablated (Fig. 1F). Crucially, there was no change in labeling for the remaining indoles supporting their classification as host derived metabolites.

Thus, the full scope of measured indole metabolites made from tryptophan includes four host derived (indole-3-pyruvate, indole-3-lactate, indole-3-acetate, and indole-3-carboxylic acid) and two microbiome-derived (indole-3-propionate and indoxyl sulfate).

### Transaminase and downstream products of phenylalanine are also host derived

Similar to indole metabolites, phenol metabolites from phenylalanine and tyrosine are considered primarily microbiome derived (15). Other than changes in the aryl side group between tryptophan, phenylalanine, and tyrosine, the enzymatic transformations needed to produce the physiologic important phenols from phenylalanine or tyrosine mimic the transformations needed to produce indoles from tryptophan. To understand if there is also host production of phenylalanine derived metabolites, we analyzed serum via LC-MS from germ-free and control mice (Fig. 2A). As with the indoles, to mitigate the effect of dietary phenols, mice were fed a purified diet for 10 days, however even this diet contained detectable phenols (phenylpyruvate, phenyllactate, and benzoic acid, Fig. S2A). The transaminase product of phenylalanine, phenylpyruvate, and its reduced product, phenyllactate, were not changed between germ-free and control mice (Fig 2B). Cinnamic acid, the alpha-beta unsaturated metabolite of phenylalanine was also not significantly changed between germ-free and control mice. Phenylacetate was below detectable levels in all the serum samples. Benzoic acid and 3-phenylpropionate were significantly decreased in germ-free mice.

**Figure 2:**
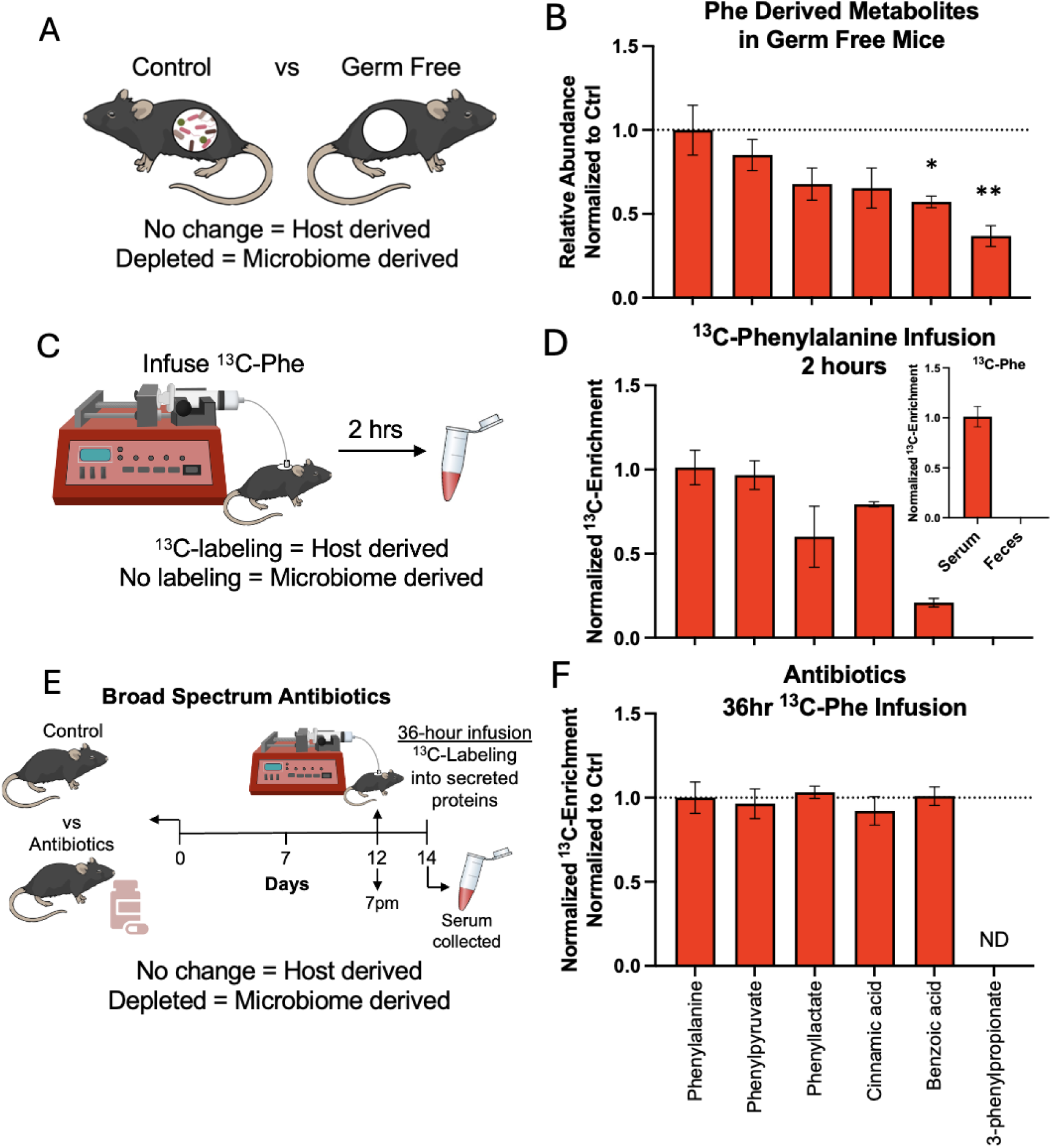
Transaminase and downstream products of phenylalanine are host derived. (**A**) Scheme and (**B**) measurement of phenylalanine derived phenol metabolites in serum of GF mice (n=5 female mice) compared to Ctrl mice (n=4 female mice). While there is no microbial contribution in GF mice, circulating metabolite concentrations may be affected by direct dietary input of the metabolites. (**C**) Scheme describing the infusion of ^13^C-phenylalanine for two hours and (**D**) ^13^C-enrichment of phenylalanine-derived metabolites in serum normalized to serum ^13^C-phenylalanine enrichment (n=3 male mice). Inset showing ^13^C-phenylalanine in serum and feces at 2 hours. (**E**) Scheme detailing a 14-day antibiotic treatment with ^13^C-phenylalanine infused for the last 36 hours. (**F**) ^13^C-phenol enrichment in antibiotic treated male mice (n=3) compared to enrichment of ^13^C-phenol metabolites in control male mice (n=3). *p* values are notated as \**p*<0.05 and \*\**p*<0.01 as calculated by a two-tailed *t*-test. ND= not detected.

Similarly to tracing indole metabolite production using ^13^C-tryptophan, we used stable isotope labeling to measure the production of the above phenols from ^13^C-phenylalanine (Fig. 2C). A minimally perturbative jugular vein infusion of ^13^C-phenylalanine for 2 h, there was no detectable ^13^C-phenylalanine in the feces. All phenylalanine-derived phenols that were not changed in GF mice (phenylpyruvate, phenyllactate, and cinnamic acid) were, however, already labeled by the infused ^13^C-phenylalnine tracer by 2 h (Fig. 2D). Phenylpyruvate was labeled quantitatively and phenyllactate and cinnamic acid enrichment reached ∼60% and 80% relative to phenylalanine. Regarding phenol metabolites that were significantly decreased in the GF mice, 3-phenylpropionate was not detectably labeled in serum 2 h into the ^13^C-phenylalalnine infusion, while benzoic acid was enriched but only by ∼20% (Fig. 2D). Based on these observations, we assigned phenylpyruvate, phenyllactate, cinnamic acid as host derived, benzoic acid as partially host derived, and 3-phenylpropionate as microbiome derived.

To achieve labeling of microbially derived phenols, we infused the ^13^C-phenylalalnine tracer for 36 hours. By 36 hours into the infusion, the circulating ^13^C-phenylalanine enrichment reached ∼25% and the fecal enrichment reached ∼7% (Fig. S2B). As with the 2-hour infusion, there was full enrichment of phenylpyruvate and ∼80% enrichment of phenyllactate and cinnamic acid (Fig. S2C). Even after 36 hours of a ^13^C-phenylalalnine infusion, benzoic acid was still only enriched by ∼30%. Interestingly, no ^13^C-3-phenylpropionate was detected. As the infused ^13^C-phenylalanine only gets to the gut microbiome through secreted protein, the missing labeling for benzoic acid and 3-phenylpropionate implies another upstream precursor for these metabolites. This could be from dietary contamination or bacterial digestion of dietary protein.

We next infused ^13^C-phenyalalnine in antibiotics treated mice for 36 hours (Fig. 2E). ^13^C-phenylalanine labeling in the circulation (∼25%) or feces (∼7%, Fig. S2C) was unaffected by antibiotics. Additionally, enrichment of all the host derived metabolites (phenylpyruvate, phenyllactate, cinnamic acid, and benzoic acid) was maintained in antibiotics-treated mice, further supporting the host origin of these metabolites (Fig. 2F).

### Host metabolism is responsible for circulating hydroxyphenylpyruvate and hydroxyphenyllactate

Tyrosine is structurally identical to phenylalanine except for a hydroxyl group on the aryl ring. As such, we hypothesized that tyrosine may also undergo the same host-mediated transformations to hydroxyphenylpyruvate and hydroxyphenyllactate. Additionally, phenol and cresol are metabolites that are understood to come from microbial digestion of tyrosine down to the aryl group or aryl group plus one remaining methyl group. These metabolites then get sulfated by the host. To understand the host contribution to tyrosine-derived metabolites, we measured these metabolites in serum from germ-free and control mice (Fig. 3A). All mice were fed a purified diet which had lower concentrations of many of the tyrosine-derived metabolites compared to chow (Fig. S2D). Surprisingly, however, the purified diets had higher concentrations of phenol sulfate and cresol sulfate than chow. Similar to the host metabolism of phenylalanine and tryptophan, there was no change in circulating hydroxyphenylpyruvate and hydroxyphenyllactate in germ-free vs control mice (Fig 3B). Hydroxyphenylacetate and hydroxycinnamic acid were not at detectable levels in any of the serum samples. Cresol sulfate and phenol sulfate were significantly depleted in germ-free mice.

**Figure 3:**
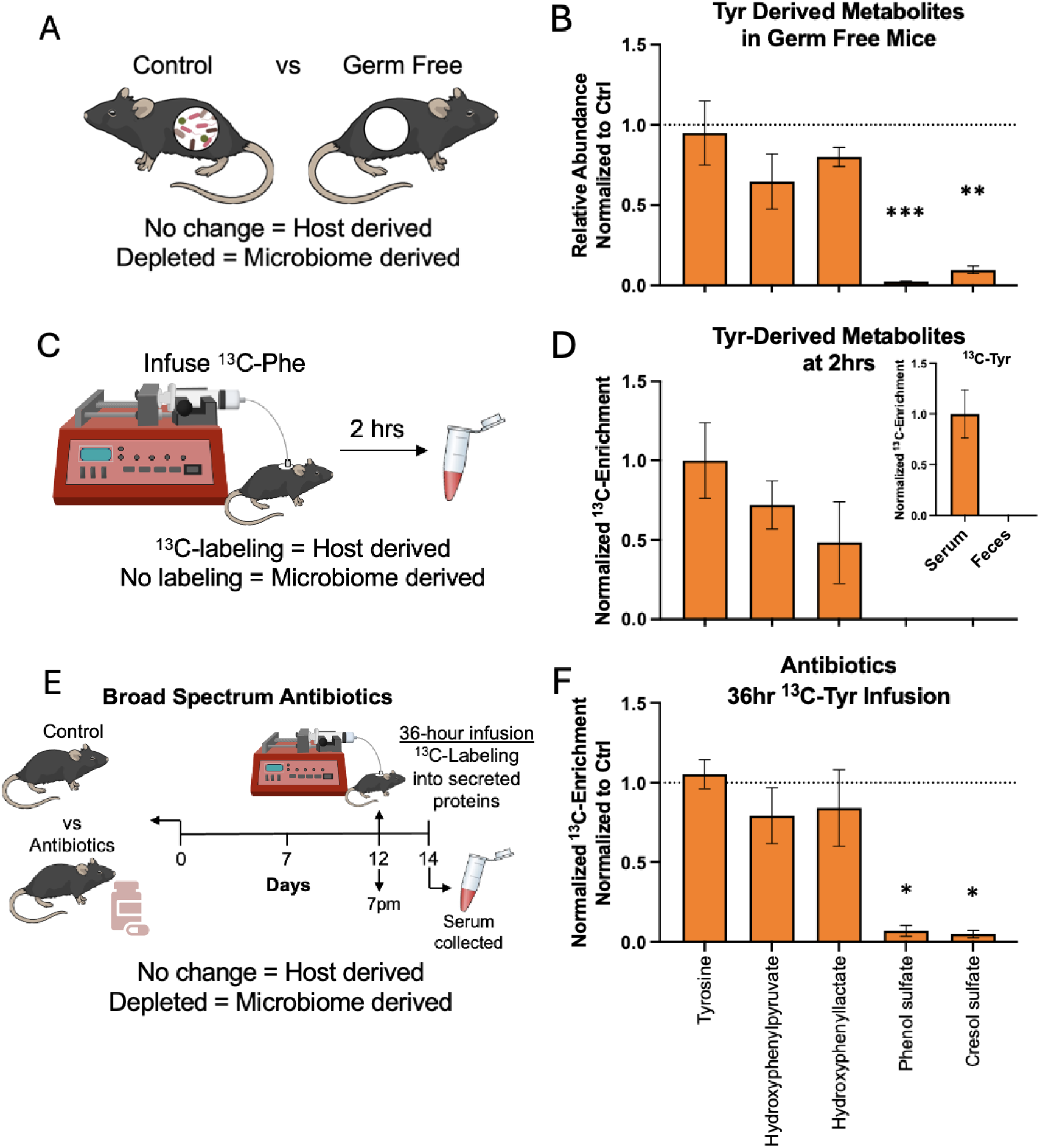
Transaminase products of tyrosine are host derived. (**A**) Scheme and (**B**) measurement of indole metabolites in serum of GF mice (n=5 female mice) compared to Ctrl mice (n=4 female mice). (**C**) Scheme describing the infusion of ^13^C-phenylalanine for two hours and (**D**) ^13^C-enrichment of tyrosine-derived metabolites in serum normalized to serum ^13^C-tyrosine enrichment from the ^13^C-phenylalanine infusion. Inset showing ^13^C-tyrosine in serum and feces at 2 hours (n=3 male mice). (**E**) Scheme detailing a 14-day antibiotic treatment with ^13^C-tyrosine infused for the last 36 hours. (**F**) ^13^C-phenol enrichment in antibiotic treated male mice (n=3) compared to enrichment of ^13^C-phenol metabolites in control male mice (n=3). \**p*<0.05, \*\**p*<0.01, and \*\*\**p*<0.001 as calculated by a two-tailed *t*-test.

We next sought to trace host production of tyrosine-derived metabolites. The insolubility of ^13^C-tyrosine in saline at neutral pH limited the concentration at which tyrosine could be infused. At 2 hours post infusion with ^13^C-tyrosine, the serum enrichment reached only ∼1% (Fig. S2E), which was too low to consistently see labeling into downstream metabolites. As tyrosine can be made from phenylalanine, we measured ^13^C-enrichment of tyrosine post an infusion of ^13^C-phenylalanine for 2 hours. The ^13^C-tyrosine enrichment from a phenylalanine infusion reached ∼2%, which was enough to reliably see ^13^C-enrichment in hydroxyphenylpyruvate and hydroxyphenyllactate (Fig. 3D). Normalizing to the tyrosine enrichment, hydroxyphenylpyruvate was enriched by 72% and hydroxyphenyllactate was enriched by 48%. No ^13^C-phenol or cresol sulfate was detected. Overall, we define hydroxyphenylpyruvate and hydroxyphenyllactate as primarily host derived, whereas phenol sulfate and cresol sulfate are microbially derived.

To achieve labeling of phenol and cresol sulfate, we infused ^13^C-tyrosine for 36 hours. By 36 hours post infusion, the circulating ^13^C-tyrosine enrichment was still low, reaching only ∼4% (Fig. S2F). Despite this, fecal ^13^C-tyrosine could be reliably detected, reaching ∼2% (Fig. S2G). All tyrosine derived metabolites were enriched by 36 h with enrichment relative to serum tyrosine reaching 75% for hydroxyphenylpyruvate, 61% for hydroxyphenyllactate, 70% for phenol sulfate, and 48% for cresol sulfate. Like benzoic acid, the remaining contribution could be from bacterial digestion of dietary protein.

To differentiate labeling by host metabolism versus microbiome metabolism, we infused ^13^C-tyrosine in antibiotics-treated mice (Fig. 3E). Consistent with our definition of hydroxyphenylpyruvate and hydroxyphenyllactate as host derived, there was no change in enrichment between control and antibiotics-treated mice for these two metabolites (Fig. 3F). There was, however, a significant decrease with antibiotics in ^13^C-enrichment of phenol sulfate and cresol sulfate, consistent with these metabolites being primarily microbially derived.

### Overall concentrations of indole and phenol metabolites are similar in feces as in circulation

As indole and phenol metabolites have been considered purely microbiome metabolites, many studies focus on their fecal concentrations. To understand if there is a difference between fecal and circulating concentrations, we measured, for mice on a purified diet, the absolute indole and phenol metabolite concentrations in serum and feces. The most abundant metabolites overall were cresol sulfate, phenylpyruvate, hydroxyphenyllactate, and indole-3-pyruvate (Fig. S3). Even though cresol sulfate production requires the microbiome (to make cresol), it is more abundant in the systemic circulation than feces as the sulfation occurs in the liver. A similar pattern is seen for indoxyl sulfate. Some products that are made solely by the microbiome such as indole-3-proprionate nevertheless showed similar fecal versus serum levels, reflecting effective sharing from the microbiome to the host. Similarly, some products that are actively produced in the host, such as hydroxyphenyllactate, nevertheless showed high fecal concentrations, likely reflecting local production also within microbes. Overall, the substantial levels of these compounds in both feces and serum, with no consistent pattern as to where they are more abundant, is consistent with many of the compounds being made by both host and microbiome.

### Human cell lines can produce indole and phenol metabolites

As many indole and phenol metabolites can be made by the host in vivo, we wanted to know if cultured mammalian cells can also autonomously produce these metabolites. To do this, we cultured colorectal cancer cells (HCT116) and traced metabolism of tryptophan, phenylalanine, and tyrosine (Fig. 4A). In cells cultured with media containing ^13^C-tryptophan, indole-3-pyruvate and indole-3-lactate were labeled by the added ^13^C-tryptophan tracer (Fig. 4B). Indole-3-pyruvate was labeled to the same quantitative extent as the tryptophan tracer. While dialyzed fetal bovine serum was used for these experiments, there was still unlabeled indole-3-lactate detected in the fresh media. This diluted the measured enrichment, but nevertheless indole-3-lactate labeling was substantial. The other host derived indole metabolites (indole-3-acetate and indole-3-carboxylic acid) were not made in HCT116 cells.

**Figure 4:**
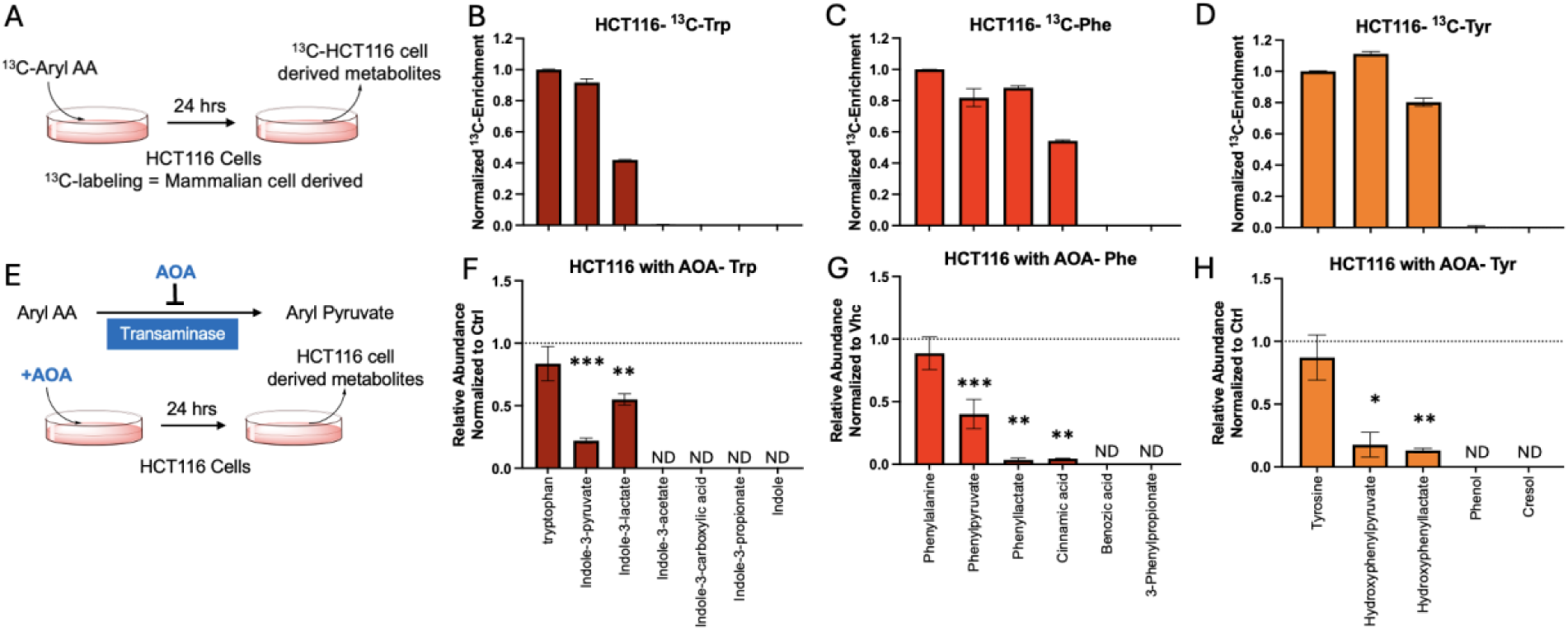
Many indoles and phenols are made in human cells *in vitro* via transaminase. (**A**) Schematic detailing tracing metabolism of ^13^C-amino acids into indoles and phenols in HCT116 cells. (**B**) ^13^C-enrichment of indole metabolites in HCT116 cells 24 hours post the addition of 16mg/mL ^13^C-tryptophan to the media. (**C-D**) ^13^C-enrichment of phenol metabolites from (C) 66mg/mL ^13^C-phenyalalnine or (D) 80mg/mL ^13^C-tyrosine. In (B-D), enrichment was normalized to the respective ^13^C-amino acids in HCT116 cells. (**E**) Schematic showing HCT116 cells being treated with 500uM AOA to inhibit transaminase, the first step for most of the indole and phenol synthesis. (**F**) Abundance of indole metabolites 24-hours post addition of AOA compared to cells treated with vehicle (Vhc). (**G-H**) Abundance of phenylalanine (G) or tyrosine (H) derived phenol metabolites 24-hours post addition of AOA. (H) compared to cells treated with vehicle. ND=not detected in vehicle or AOA treated cells. \**p*<0.05, \*\**p*<0.01, and \*\*\**p*<0.001 as calculated by a two-tailed *t*-test.

Similarly, we cultured HCT116 cells in media containing either ^13^C-phenylalanine or ^13^C-tyrosine. Consistent with their being host-derived metabolites, the cells produced phenylpyruvate, phenyllactate, and cinnamic acid from the added ^13^C-phenylalanine. The microbiome metabolite 3-phenylpropionate was, as expected, not made from the added tracer (Fig. 4C). Benzoic acid, which comes at least partially from host metabolism in mice, was not produced in HCT116 cells. From ^13^C-tyrosine, HCT116 cells were also able to produce labeled hydroxyphenylpyruvate and hydroxyphenyllactate with no production of phenol or cresol (Fig. 4D). Because many of these metabolites have classically been considered microbial products, we confirmed the sterility of our HCT116 cultures. The HCT116 cells were confirmed to be negative for *Mycoplasma* contamination (Fig. S4). Production of these metabolites was also observed in various other common cell lines (Fig. S5). Overall, cultured mammalian cells had similar metabolic capacity to the host *in vivo,* converting all aryl-amino acids to their respective aryl-pyruvates, aryl-lactates, and cinnamic acid.

Metabolism from aryl amino acids to the indole and phenol metabolites could logically proceed initially through transamination to the respective pyruvate analogues. To assess whether transaminase inhibition could block production of the downstream products, we treated HCT116 cells with the pan-transaminase inhibitor, aminooxyacetic acid (AOA, Fig. 4E). AOA was cytostatic at the given dose (500uM). Upon treatment with AOA, all indole and phenol metabolites made by HCT116 cells (indoles: indole-3-pyruvate and indole-3-lactate; phenols: phenylpyruvate, phenyllactate, cinnamic acid, hydroxyphenylpyruvate, and hydroxyphenyllactate) were depleted (Fig. 4F-H). Thus, isolated mammalian cells can autonomously produce many indole and phenol metabolites through transamination followed by subsequent metabolic reactions.

### IL4i1 is not the main host contributor to indole-3-pyruvate or phenylpyruvate

Recently, IL4i1, an enzyme initially discovered as being induced by IL4 in B cells, but since found to be expressed and excreted predominately by monocyte lineage cells, has been characterized to convert phenylalanine to phenylpyruvate and tryptophan to indole-3-pyruvate *in vitro* (19–21). To assess contribution of IL4i1 to overall host indole and phenol pools, we analyzed plasma from IL4i1-/- mice (Fig. S6A). There were no significant changes in any of the host derived indole metabolites between wild type and IL4i1-/- C57BL6 mice (Fig. S6B). Additionally, there were no significant differences between IL4i1-/- and wild type mice of any of the host-derived metabolites from tyrosine or phenylalanine (Fig. S6C-D).

### Characterization of the reactions in lysates

To further characterize the mammalian synthetic pathway to indoles and phenols, we created lysates of HCT116 cells (Fig. S7A). The conversion of the amino acid to aryl-pyruvate is likely through a transaminase which typically require the cofactor pyridoxal 5’-phosphate (PLP) as well as a nitrogen acceptor such as alpha-ketoglutarate (*α*KG). To assess the transamination step, we added phenylalanine and tryptophan as well as PLP and *α*KG to the lysates. In the HCT116 lysates with PLP and *α*KG, there was robust conversion of tryptophan to indole-3-pyruvate reaching ∼55% conversion by 4 hours (Fig. S7B). There was no conversion without the addition of PLP and *α*KG. While there was production of phenylpyruvate in the HCT116 lysates, the reaction stalled by 15 minutes and the maximum yield was <1% (Fig. S7C). This suggests the lysate contains this enzymatic activity, but the enzyme is unstable under the tested reaction conditions.

Additionally, conversion of the pyruvate species to the aryl-lactates likely requires a reducing agent like NADH. The addition of NADH with indole-3-pyruvate and phenylpyruvate to the lysates allowed for the production of indole-3-lactate (17% yield) and phenyllactate (12% yield) by 4 hours (Fig. S7D-E). Furthermore, cinnamic acid was also produced from phenylpyruvate with the addition of NADH (3%). Interestingly, when phenyllactate was added with NADH in the lysates (Fig. S7F), no cinnamic acid was formed, suggesting cinnamic acid production, at least in HCT116 cells, may be directly from phenylpyruvate.

### Patients treated with antibiotics have no change in host-derived indole and phenol metabolites

The above data clearly differentiate indole and phenol metabolites into host-derived and microbiome-derived metabolites. We next asked if this categorization would align with metabolic responses in human patients receiving antibiotics. To this end, we analyzed the abundance of indole and phenol metabolites in a publicly available metabolomics data set of serum from patients in general admissions at Boston Children’s Hospital (25, study MTBLS948). The original purpose of this dataset was to compare metabolic profiles of generally healthy individuals (ages 2 months to 55 years old) against a cohort of pediatric patients with a variety of diseases including rare metabolic or genetic diseases. Although the intention was not to study the effect of antibiotics, many patients had detectable levels of antibiotics in their serum. Out of the 169 patients in the data set, 16 had detectable circulating levels of at least one of the following antibiotics: ciprofloxin, moxifloxin, clindamycin, erythromycin, sulfociprofloxacin, doxycycline, metronidazole, sulfamethoxazole.

When comparing serum from patients who had detectable levels of antibiotics to those without, there was no change in tryptophan-derived host metabolites (indole-3-lactate, indole-3-acetate, and indole-3-carboxylic acid, Fig. 5A). For microbial tryptophan-derived metabolites, indole-3-propionate was significantly decreased in patients with antibiotics (Fig. 5B). While indoxyl sulfate was not changed for patients receiving a single antibiotic, there was a significant decrease in indoxyl sulfate for patients receiving more than one antibiotic (p<0.0001).

**Figure 5:**
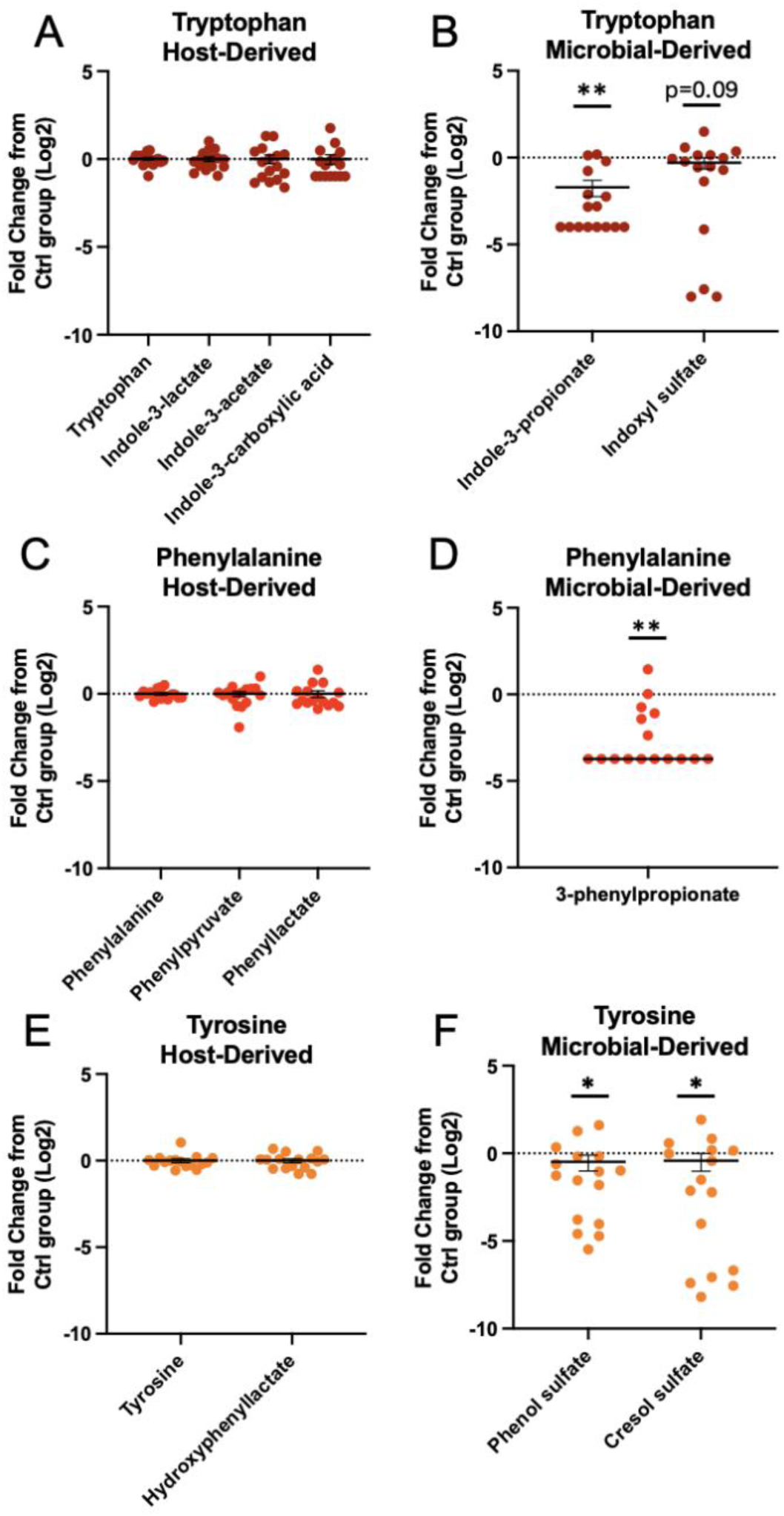
Human aryl-amino acid metabolism to indoles and phenols mirrors that of mice. (**A-B**) Log2 fold change of (**A**) host derived and (**B**) microbial derived serum indole metabolites in patients with (n = 16) or without (n=153) detectable levels of antibiotics in circulation. (**C-F**) Fold change of (**C**) host synthesized or (**D**) microbial synthesized phenols derived from phenylalanine, and (**E**) host synthesized or (**F**) microbial synthesized phenols derived from tyrosine. *p* values are notated as \**p*<0.05 and \*\**p*<0.01 as calculated by a two-tailed *t*-test.

Additionally, phenylalanine-derived host metabolites (phenylpyruvate and phenyllactate) did not change between patients on antibiotics and the control group (Fig 5C), but microbially derived 3-phenylpropionate was decreased (Fig. 5D). Similarly, hydroxyphenyllactate (host derived from tyrosine) was not changed (Fig, 5E) while both phenol sulfate and cresol sulfate (microbially derived from tyrosine) were decreased in patients on antibiotics (Fig. 5F). This dataset did not include levels of the other metabolites of interest. Overall, the data suggests there is a host metabolic pathway which converts aryl-amino acids into indole and phenol metabolites that is conserved from mouse to human.

## Discussion

In this study we demonstrate that many physiologically important indole and phenol metabolites are derived from host metabolism in mice, humans and tissue culture of mammalian cells. We show there is a conserved metabolic pathway for all three aryl amino acids in mammalian cells that can convert tryptophan, phenylalanine, and tyrosine to their respective aryl-pyruvates and aryl-lactates (Scheme 1). The production of these metabolites occurs in mice and in cells in culture as shown by isotope tracing, and in humans as their concentrations are not affected by antibiotics.

Additionally, there is a mammalian metabolic pathway that can convert phenylalanine all the way to cinnamic acid. As shown in HCT116 lysates, cinnamic acid is made preferentially from phenylpyruvate directly as there was no production of cinnamic acid directly from phenyllactate. While there was no production of benzoic acid in cells in culture, our isotope tracing studies show benzoic acid production from phenylalanine is at least partially from host metabolism. There is no mammalian enzyme known to complete this conversion, but benzoic acid could feasibly be made from cinnamic acid through a mechanism similar to the cinnamate:CoA ligase enzyme as seen in plant peroxisomes (26).

Further, host metabolism is responsible for the production of indole-3-acetate and indole-3-carboxylic acid from tryptophan. While the mammalian enzymes are not known, indole-3-acetate is known to be produced from indole-3-pyruvate in bacteria and plants (27,28). The exact pathway for indole-3-carboxylic acid production in bacteria is not clear (29), but the conversion of indole-3-acrylic acid to indole-3-carboxylic acid mimics the conversion of cinnamic acid to benzoic acid.

Not all indole and phenol metabolites can be made through host metabolic pathways. For example, bacterial metabolism is required for freeing the aryl side group into indole, phenol, or cresol. Additionally, the gut microbiome is necessary to reducing the amino acid all the way down to indole-3-propionate or 3-phenylpropionate.

Indole-3-lactate and indole-3-acetate are frequently studied metabolites as they convey important anti-inflammatory affects, with supplementation protecting in mouse models from diseases ranging from inflammatory bowel disease to extreme nearsightedness (2,7,30,31). Often changes in these indole metabolite concentrations are correlated to observed microbiome composition changes in patient populations or mouse models. These correlations are classically interpreted as evidence that microbiome metabolism is responsible for metabolite production. Based on our data demonstrating indole metabolites are predominately host derived, we propose three potential alternative explanations to these disease-microbiome-metabolite correlations: (i) the disease state could affect microbial composition and host production of indole metabolites independently, (ii) the effect of the disease on host indole production could affect microbiome composition, or (iii) microbial changes from the disease could be affecting host metabolic pathways.

Overall, an understanding of the source of indole and phenol metabolites is the fundamental first step to developing tools to control their circulating concentrations. Here we show that host metabolism, rather than microbiome composition, is a primary source for physiologically important metabolites such as indole-3-lactate, indole-3-acetate and cinnamic acid.

## Experimental Models

### Mouse studies

All mouse studies were completed in compliance with protocols approved by the Princeton University Animal Care and Use Committee. Unless otherwise indicated, all mice were C57BL/6NCrl mice (Charles River Laboratories, strain 027) at 8-12 weeks old and fed a standard purified diet (Research Diets Inc, D11112201). Mice were group housed on a normal light-dark cycle (8:00-20:00) and had ad lib access to food and water.

### Germ free mouse model

C57BL/6N wild type female germ-free or murine pathogen free mice (Taconic strain B6-F GF or MPF) were housed in individually vented cages (Allentown Sentry Sealed Positive Pressure system) under a 12-hour light and 12-hour dark cycle. All mice were 9-10 weeks old, group housed and had free access to water and doubly irradiated standard purified diet (D11112201-1.5Vii). Contamination quality checks were performed on mouse fecal pellets, cage samples, and research diet pellets via aerobic and anaerobic cultures. After 10 days on the diet, serum and fecal samples were collected, and metabolomics analysis was completed via liquid chromatography-mass spectrometry (LC-MS).

## Methods

### In vivo sample collection

Serum samples were collected via tail bleed. Blood was placed on ice immediately following collection, centrifuged at 4 °C, 14,000xg for 10 min, and moved to another tube to be stored at −80 °C. Feces were collected fresh, immediately placed on dry ice and stored at -80°C until processing.

### Metabolite extraction

Serum samples were extracted with methanol (3uL serum into 120uL methanol) and incubated for 10 minutes on ice. The samples were then centrifuged at 15,000xg for 30 min at 4°C and the resulting supernatant was transferred to a mass spectrometry vial for further analysis. Fecal samples were collected fresh and immediately placed on dry ice. Fecal and dietary samples were ground at liquid nitrogen temperature with a cryomill (Restch, Newtown, PA). The resulting powder was extracted with 40:40:20 methanol: acetonitrile: water (40 ul extraction solvent per 1 mg tissue) for 10 min on ice and centrifuged at 15,000xg for 10 min. The resulting extract was then moved to LC-MS vials (Thermo Fisher Scientific, 200046; caps, Thermo Fisher Scientific, 501313) for measurement.

### Metabolite measurement

For measurement of water-soluble metabolites, a Q Exactive Plus hybrid quadrupole-orbitrap mass spectrometer (Thermo Fisher Scientific) coupled with hydrophilic interaction chromatography (HILIC). For LC, an An XBridge BEH Amide column (150 mm × 2.1 mm, 2.5 μM particle size, Waters) was used with a gradient solvent A (95%:5% H_2_O: acetonitrile with 20 mM ammonium acetate, 20 mM ammonium hydroxide, pH 9.4) and solvent B (100% acetonitrile). The gradient ran as follows: 0 min, 90% B; 2 min, 90% B; 3 min, 75% B; 7 min, 75% B; 8 min, 70% B, 9 min, 70% B; 10 min, 50% B; 12 min, 50% B; 13 min, 25% B; 14 min, 25% B; 16 min, 0% B, 20.5 min, 0% B; 21 min, 90% B; 25 min, 90% B with a flow rate of 150 μl min^−1^. 10 μL was used as the injection volume and the column temperature was set to 25°C. A negative-ion mode was used for the MS scans with a resolution of 140,000 at m/z 200. The automatic gain control target was 5 × 10^6^ and the scan range was *m*/*z* 75−1,000. Xcalibur (v.4.3; Thermo Fisher Scientific) was used to collect raw data.

All LC-MS data was analyzed using El-Maven (v.0.12.1)(32). All ^13^C-labeling data was corrected for natural ^13^C abundance using the Accucor package (33) (https://github.com/XiaoyangSu/AccuCor).

### ^13^C-amino acid compounds

All amino acids were from Cambridge isotopes (^13^C_9_-Phenylalanine-CLM-2250-H, ^13^C_9_-Tyrosine-CNLM-439-H and ^13^C_11_-Tryptophan-CLM-4290-H). Upon suspicion of contamination of the ^13^C-tryptophan with ^13^C-indole-3-acetate, the ^13^C-tryptophan was further purified by Lotus Separations, LLC using a Phenomenx Luna® Amino column (100 Å, 5 µm, 150 (L) x 21 (ID) mm). Separation conditions were as follows: isocratic conditions with 60% A (*20 mM ammonium acetate, 20 mM ammonium hydroxide, 95:5 water:acetonitrile, pH 9.6), and 40% B (acetonitrile) at 45°C* and a flow rate of 15mL min^−1^.

### Jugular vein and carotid artery catheterization

For jugular vein catheterized mice, using aseptic surgical techniques, a vascular access button (Instech, VABM1B/25, 25 gauge, one-channel button) was implanted under the back skin of a mouse, and connected to a catheter (Instech, C20PV-MJV1301, 2 French, 10 cm) that was placed in the right jugular vein. Mice were allowed at least 5 days to recover from surgery before any tracer infusion. Catheters were flushed with sterile saline and refilled with sterile heparin glycerol locking solution (SAI, HGS) at least once a week.

### 2-hour ^13^C-amino acid infusions

All infusions were completed with the mice fed ad libitum. Mice were weighed to calculate the tracer infusion rate. At 18:00, mice were attached to the infusion line with a swivel and tether (Instech, swivel, SMCLA; line, KVABM1T/25) and allowed to acclimate for one hour. With an infusion pump (SyringePump, NE-1000), the infusate was advanced through tubing into the mouse catheter at a rate of 0.1 μL min^−1^ per gram mouse weight starting at 19:00. Infusates used were 5mM ^13^C-tryptophan, 2mM ^13^C-tyrosine, or 15mM ^13^C-phenylalanine in sterile saline. After two hours of infusion, serum was collected by tail snip and feces were collected fresh and immediately placed on dry ice. All collected samples were stored at -80°C prior to processing.

### 36-hour infusion with and without antibiotic treatment

Antibiotics were administered through addition to the mouse drinking water. An antibiotic cocktail including 1 g/L ampicillin, 1 g/L neomycin, 1 g/L metronidazole, and 1 g/L vancomycin, and 0.5% aspartame was added to make the water more palatable. Infusion of the ^13^C-tracers (5mM ^13^C-tryptophan, 2mM ^13^C-tyrosine, 15mM ^13^C-phenylalanine) was initiated on the 12^th^ day of antibiotics at 19:00 as described above. Serum and feces were collected prior to the start of the infusion, 2-, 12-, and 36-hours post initiation of the infusion. Mice had free access to food throughout the infusion.

### Cell culture

All cells obtained from ATCC, and stock cells were maintained in either DMEM (HCT116, A549, HepG2, and Hek293) or RPMI (THP-1 and Jurkat) with 10% complete (nondialyzed) FBS. All cell lines were cultured at 37°C and 5% CO2. Cells were grown to approximately 70%–80% confluence. Comparable cell density was confirmed visually and by cell count with a Countess II Hemocytometer. Cells were routinely tested for *Mycoplasma* contamination using the PlasmoTest *Mycoplasma* Detection Kit (Invivogen).

### ^13^C-tracing in cell culture

All cell lines were plated into 6 well plates. Twenty-four hours post plating, the media was aspirated and switched to media containing 10% dialyzed fetal bovine serum and one of the ^13^C-aryl amino acids added at equimolar to the media concentration for that amino acid. For HCT116, A549, HepG2, and Hek293 cells, DMEM with 16mg/mL ^13^C-tryptophan, 66mg/mL ^13^C-phenylalaine, or 80mg/mL ^13^C-tyrosine added was used. For THP-1 and Jurkat cells, RPMI with 5mg/mL ^13^C-tryptophan, 15mg/mL ^13^C-phenylalaine, or 29mg/mL ^13^C-tyrosine added was used. Cells were collected via scraping in 500uL ice cold 80% methanol post aspiration of the media. To inhibit transaminases, Aminooxyacetic acid (Sigma, C13408) was dissolved into sterile water. HCT116 cells were treated with 500uM AOA or sterile water in DMEM with dialyzed media for 24 hours.

### IL4i-/- plasma samples

Plasma samples from three female (22-23 weeks) and two male (16 weeks) whole body Il4i1 knock out mice and three female (22-23 weeks) and two male (16 weeks) wild type mice were kindly provided by Christiane Opitz (19) at the German Cancer Research Center (DKFZ), DKTK Brain Cancer Metabolism Group, 69120 Heidelberg, Germany. All mice were ad lib fed irradiated extruded chow pellets (KLIBA #3437), and at 14 days of age they also began received autoclaved oat flakes. Plasma samples were stored at -80°C and extracted and prepared for MS analysis in the same manner as serum samples.

### Cell Lysates

HCT116 cells at 50-70% confluence were trypsinized, pelleted and washed twice in ice-cold PBS before resuspending in lysis buffer (150mM NaCl, 20mM Tris-HCl at ph8, 1mM beta-mercaptoethanol, and 1X complete mini protease inhibitor cocktail with ethylene-diaminetetraacetic acid (Thermo Fischer 78429)). The pelleted cells were lysed with an Emulsiflex (Avestin). The lysed cells were spun for 60 min at 4°C at 10K rpm and allocated into 100uL aliquots with 20% glycerol and snap frozen with liquid nitrogen.

### HCT116 lysate assays

Transaminase activity was assessed in the HCT116 lysates through the addition of either phenylalanine or tryptophan (2mM) with PLP (200uM) and excess aKG (20mM). Reductase activity was assessed through the addition of indole-3-pyruvate, phenylpyruvate, or phenyllactate (2mM) and NADH (20mM). The reaction mixtures were placed at 37°C and aliquots were taken at time zero, 15 min, 1h and 4h. Reactions were run in duplicate and metabolite concentrations were quantified by LC-MS using an external standard curve.

### Human Data set

Patient sample data was obtained from a study completed at Boston Childrens’ Hospital (24). The metabolomics data set from this study was deposited to MetaboLights (24,25). A missing value imputation was performed, allowing for samples where the metabolite was below detectable limits to be filled with the minimally detected ion count for that metabolite. Out of this dataset, patients with antibiotics were selected based on detectable levels of antibiotics in circulation.

### Statistical Analysis, and Figure Production

P < 0.05 was used to determine statistical significance using a two-tailed, unpaired student’s t-test. Figures were made with GraphPad Prism (v.10.2.3), ChemDraw (v.23.1.1), and NIH BioArt (https://bioart.niaid.nih.gov/).

## Acknowledgements

Thank you to Jimmy Pratas at Lotus Separations, LLC for purifying the ^13^C-tryptophan tracer. J.E.A. is funded by F30DK139739 and M.S.D. by R01AI172144, both from the National Institute of Health. J.D.R. was supported by Ludwig Cancer Research, DP1DK113643 from the National Institute of Diabetes, Digestion and Kidney Disease, and an internal grant from the Princeton Alliance for Collaborative Research and Innovation, Princeton University. We are thankful to all members of the Rabinowitz lab for their scientific discussion and insightful input.

## Author contributions

J.E.A. and J.D.R. conceived and designed the study. J.E.A. performed the experiments, completed data analysis, and made the figures. K.O. and S.Y. assisted with in vitro assays and germ-free mouse experiments, respectively. M.R.M. performed data analysis on the patient database. C.J.H. performed the catheterization surgeries. R.P.R performed the *Mycoplasma* testing. A.H. and C.A.O. provided plasma samples from IL4i1-/- mice. M.S.D. provided crucial insights for project conceptualization. J.E.A. and J.D.R. wrote the manuscript. All authors reviewed the manuscript.

---

**Scheme 1:**
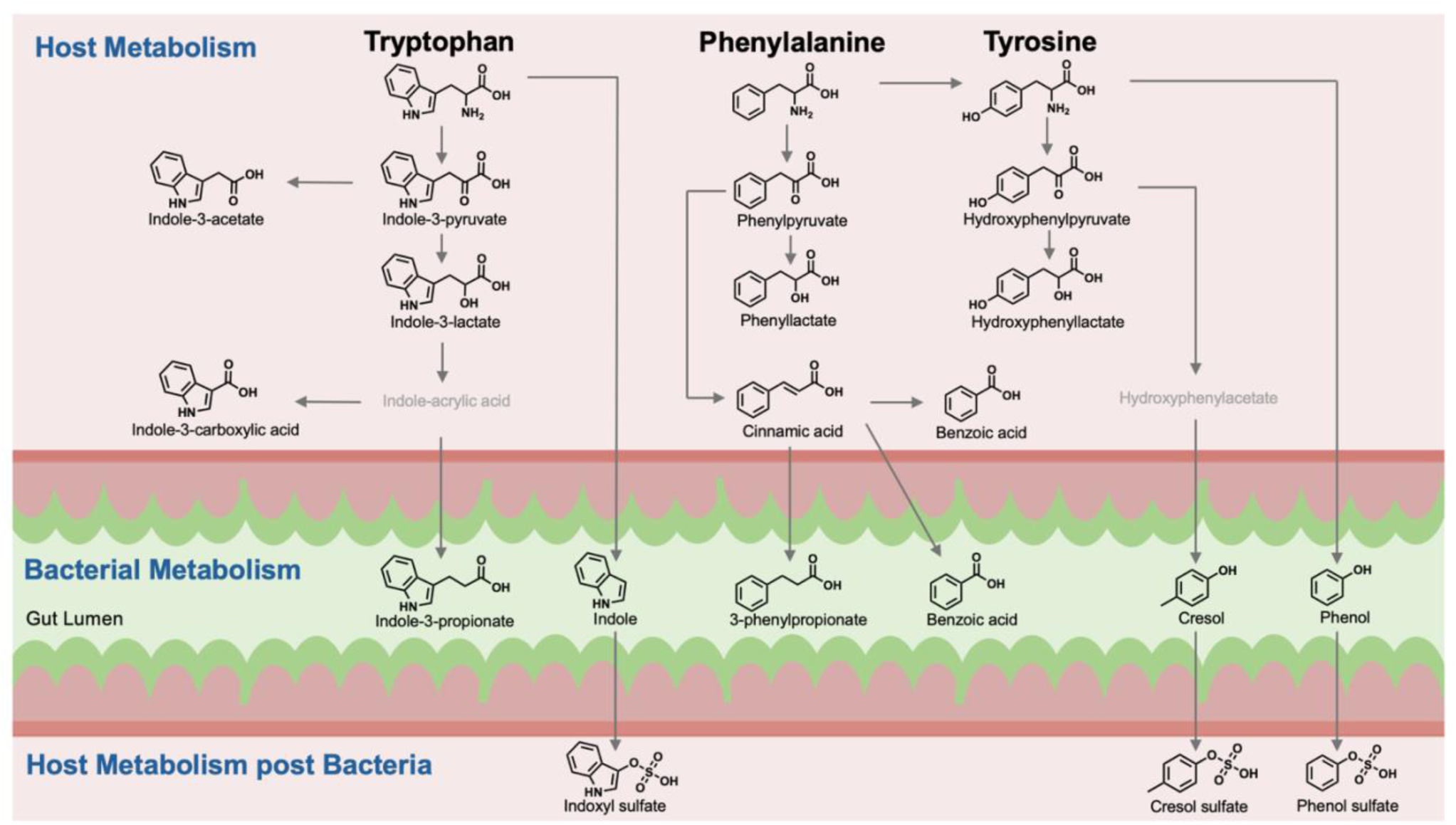
Host metabolism of aryl amino acids to indole and phenol metabolites. There is a conserved mammalian pathway across all aryl-amino acids where host enzymes can transaminate tryptophan, phenylalanine and tyrosine to their respective aryl-pyruvate and reduce to aryl-lactate. Additionally, indole-3-acetate, indole-3-carboxylic acid, and cinnamic acid are produced by host metabolism *in vivo*. While benzoic acid can also be made through host metabolism, it is still primarily a microbially derived metabolites isotope tracing of host metabolic pathways could only account for ∼30% of benzoic acid production. Bacterial metabolism is required for circulating aryl-propionates, indole, phenol, cresol.

**Figure S1:**
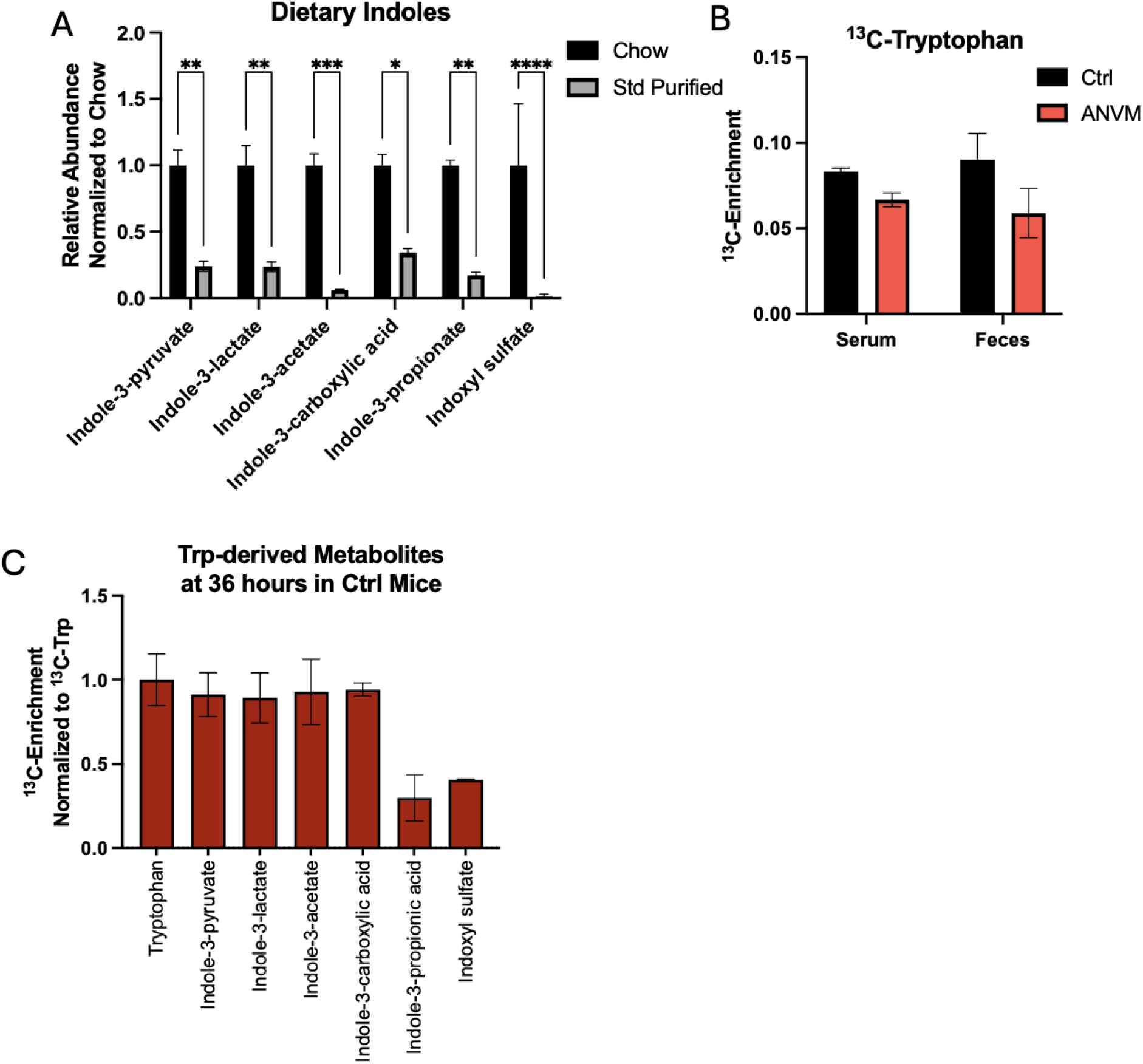
Assessing host contribution to indole metabolites. (**A**) Indole metabolites in PicoLab Chow (n=3) vs a standard purified diet (n=3). (**B**) ^13^C-enrichment of tryptophan in serum and feces after a 36-hour infusion in antibiotic treated (n=4 serum and feces) or control male mice (n=5 serum, n=3 feces). (**C**) ^13^C-enrichment of indoles in serum at the end of a 36-hour infusion of ^13^C-tryptophan in control male mice (n=3), which allows for microbially derived metabolites to be enriched. Enrichment was normalized to serum ^13^C-tryptophan enrichment. This data is used as the comparison to antibiotics-treated (ANVM) animals as seen in Figure 1F. \**p*<0.05, \*\**p*<0.01, and \*\*\**p*<0.001 as calculated by a two-tailed *t*-test.

**Figure S2:**
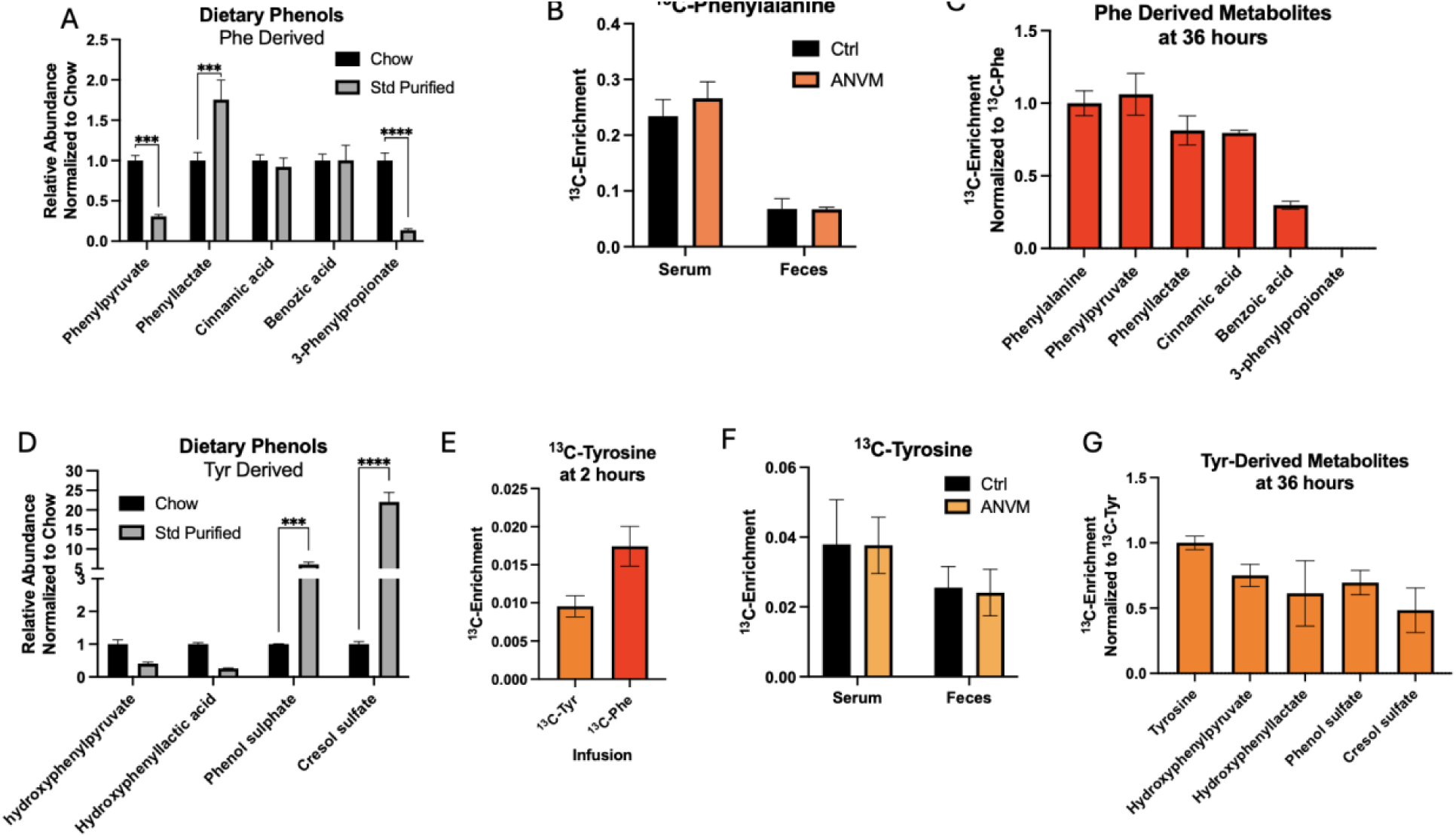
Assessing host contribution to phenol metabolites. (**A**) Phenylalanine-derived metabolites in PicoLab Chow (n=3) vs a standard purified diet (n=3). (**B**) ^13^C-enrichment of phenylalanine in serum and feces after a 36-hour infusion in antibiotic treated (n=3) or control male mice (n=3). (**C**) ^13^C-enrichment of phenols in serum at the end of a 36-hour infusion of ^13^C-phenylalanine in control male mice (n=3). Enrichment was normalized to serum ^13^C-phenylalanine enrichment. This data is used as the comparison to ANVM treated animals as seen in Figure 2F. (**D**) Tyrosine-derived metabolites in PicoLab Chow (n=3) vs a standard purified diet (n=3). (**E**) ^13^C-tyrosine enrichment in male mice infused either ^13^C-tyrosine (n=3) or ^13^C-phenylalanine (n=3) at 2 hours. (**F**) ^13^C-tyrosine in serum and feces after a 36-hour infusion in antibiotic treated (n=3) or control male mice (n=3). (**G**) ^13^C-enrichment of phenols in serum at the end of a 36-hour infusion of ^13^C-tyrosine in control male mice (n=3). Enrichment was normalized to serum ^13^C-tyrosine enrichment. This data is used as the comparison to antibiotics-treated (ANVM) animals as seen in Figure 3F. \**p*<0.05, \*\**p*<0.01, \*\*\**p*<0.001, and \*\*\*\**p*<0.0001 as calculated by a two-tailed *t*-test.

**Figure S3:**
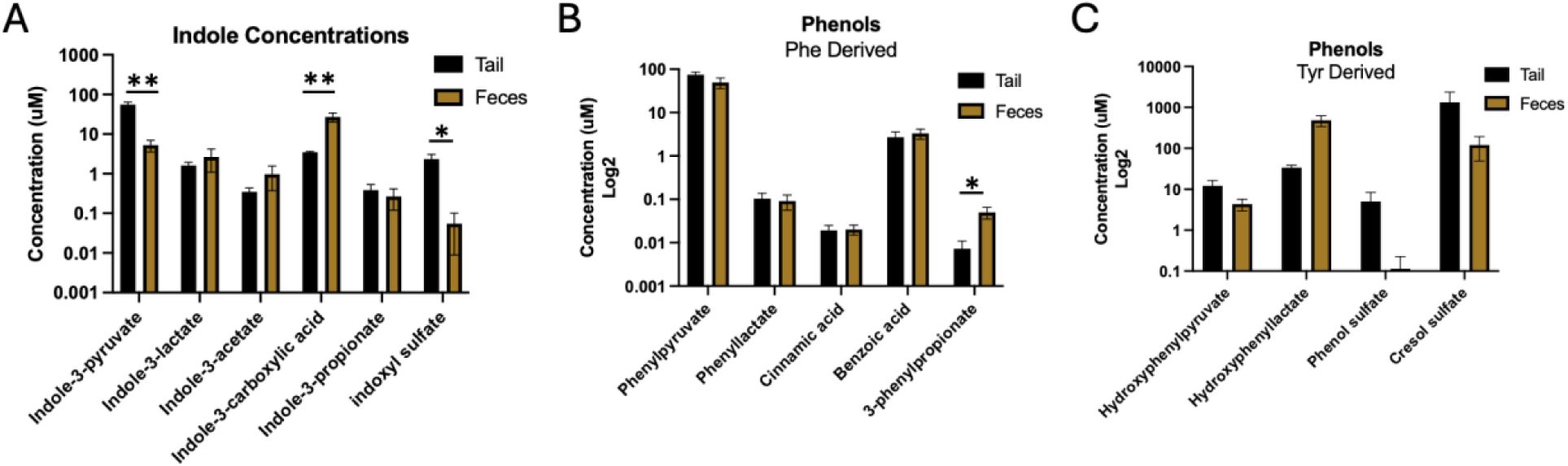
Serum and fecal concentrations of indole and phenol metabolites are comparable. **(A)** Indole metabolite concentrations in serum and feces of male mice fed a control purified diet. **(B)** Phenylalanine derived and **(C)** Tyrosine derived phenol metabolites in serum and feces of male mice fed a control purified diet. For all metabolites, n=5 serum samples, n=4 fecal samples. *p* values notated as \**p*<0.05 and \*\**p*<0.01 as calculated by a two-tailed *t*-test.

**Figure S4:**
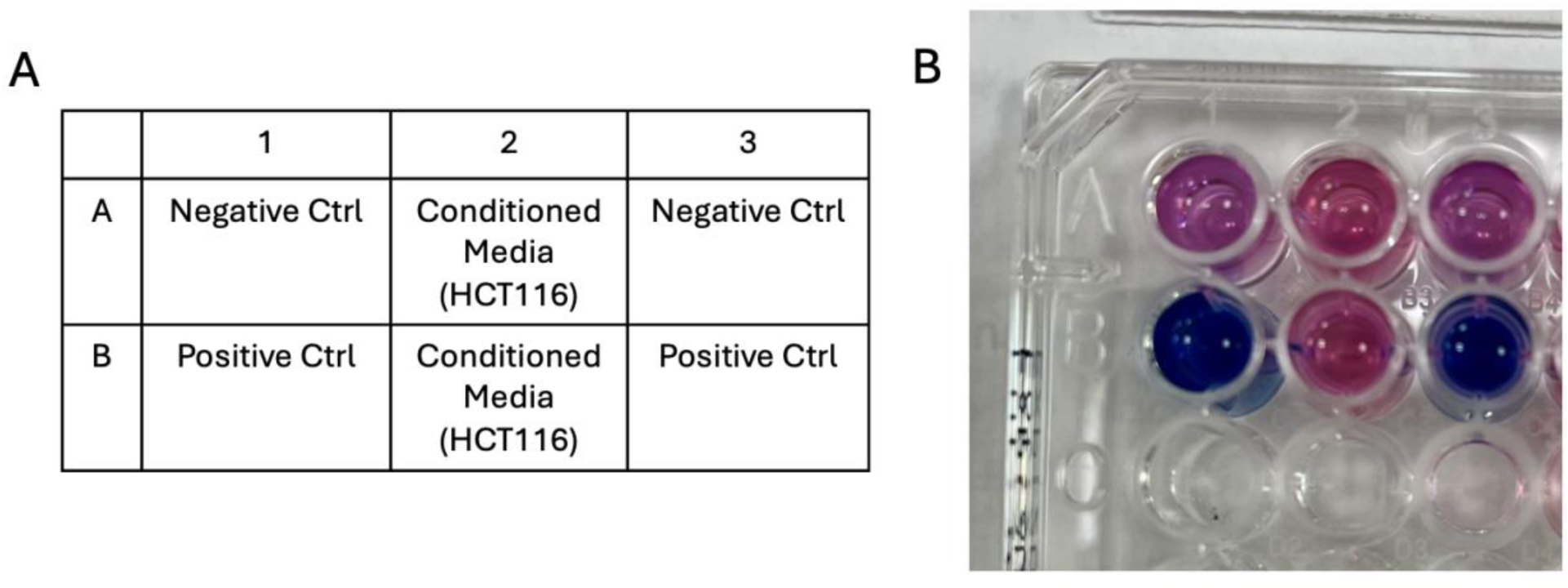
HCT116 cells were negative for *Mycoplasma* contamination. (**A**) Plate layout for the PlasmoTest *Mycoplasma* Detection Kit. (**B**) Results showing no *Mycoplasma* contamination of the HCT116 cells.

**Figure S5:**
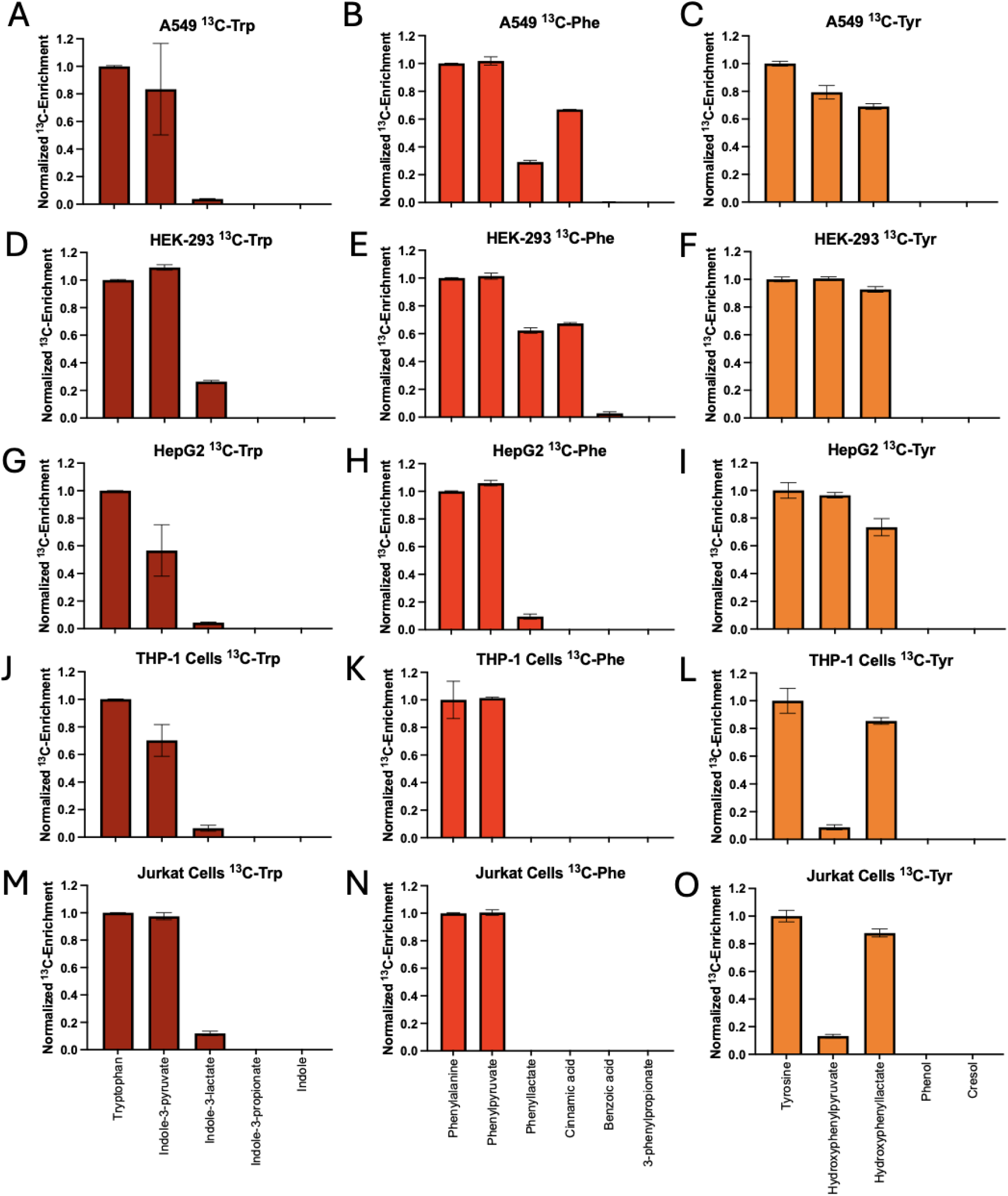
Synthesis of indoles and phenols is conserved in many different cell lines. Enrichment of indole, phenylalanine derived, and tyrosine derived metabolites in cells treated with ^13^C-tryptophan, ^13^C-phenylalalnine, or ^13^C-tyrosine for 20 hours in (**A-C**) A549 lung cancer cells, (**D-F**) HepG2 liver cancer cells, (**G-I**) HEK-293 immortalized human embryonic kidney cells, (**J-L**) THP-1 leukemia cells, and (**M-O**) Jurkat T cell lymphoma cells. All metabolite enrichment was normalized to their respective ^13^C-amino acid enrichment. All measurements are represented as the mean ± SEM with an n=3 biological replicates.

**Figure S6:**
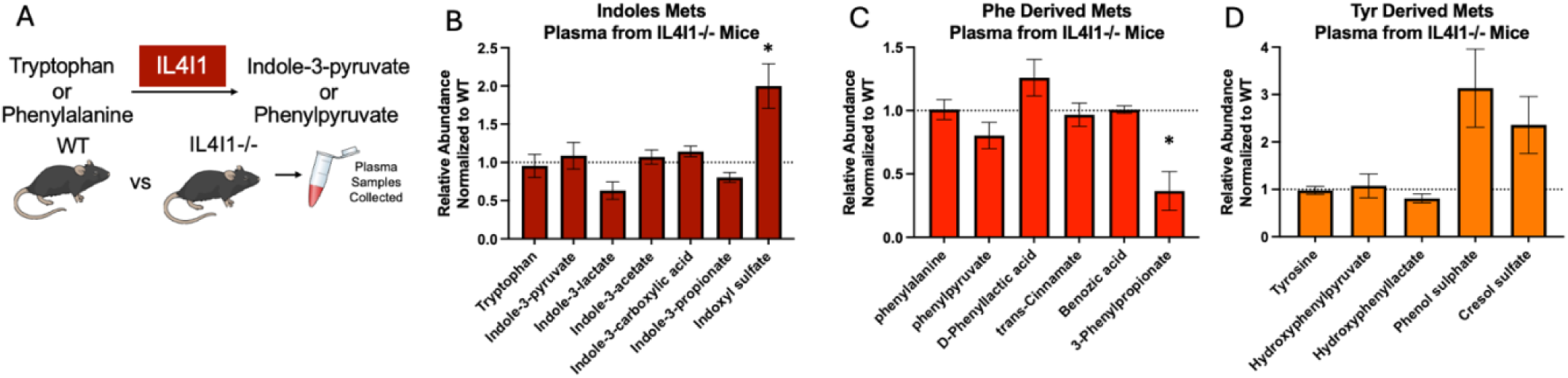
IL4I1 is not the main contributor to indole or phenol circulating metabolites. (**A**) Schematic showing the metabolic reactions of IL4I1 and the comparison of plasma from IL4I1 -/- vs wild type (WT) mice. (**B**) Indole, (**C**) phenylalanine derived phenol, and (**D**) tyrosine derived phenol metabolite abundances in whole body IL4I1-/- mice (n=3 female and 2 male mice) compared to WT mice (n=3 female and 2 male mice). \**p*<0.05 as calculated by a two-tailed *t*-test.

**Figure S7:**
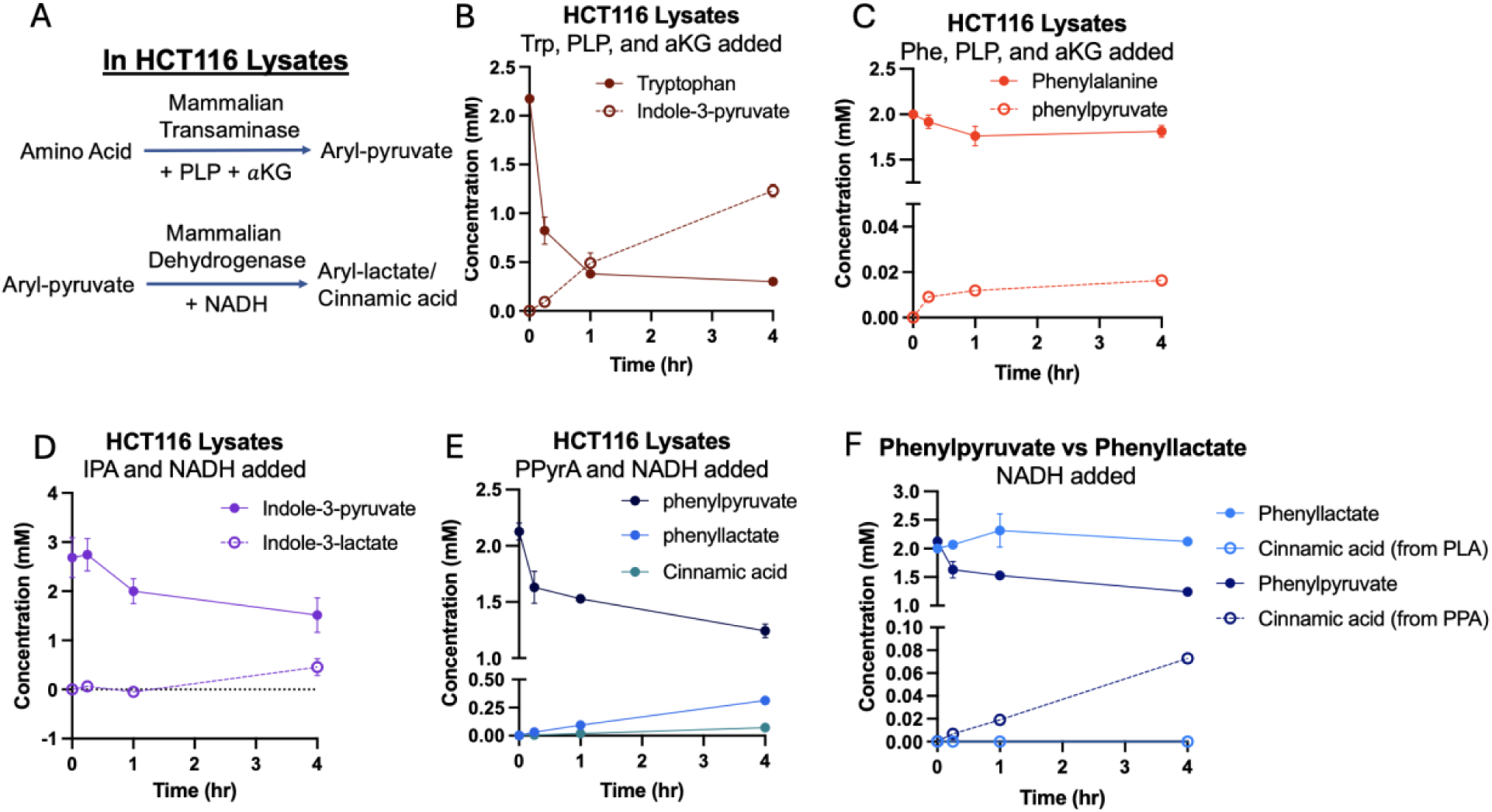
Aryl-amino acid transaminase and dehydrogenase activities are present in mammalian cell lysates. (**A**) Scheme describing the main reactions and cofactors needed in HCT116 lysates. (**B-C**) The conversion of tryptophan to indole-3-pyruvate (B) and phenylalanine to phenylpyruvate (C) occurs with the addition of pyridoxal phosphate (PLP) and alpha-ketoglutarate (*α*KG) in HCT116 lysates. No reaction occurred in lysates without the addition of PLP and *α*KG. (**D-E**) The production of indole-3-lactate (D) from the addition of indole-3-pyruvate (IPyrA) and phenyllactate, and cinnamic acid (E) from phenylpyruvate (PPyrA) with NADH in HCT116 lysates. (**F**) Cinnamic acid production from phenylpyruvate or phenyllactate and NADH in the HCT116 lysates. All measurements were performed with an n=2 independent lysate reactions.

